# Expression-based analyses indicate a central role for hypoxia in driving tumor plasticity through microenvironment remodeling and chromosomal instability

**DOI:** 10.1101/333633

**Authors:** Anqi Jing, Frederick S. Vizeacoumar, Sreejit Parameswaran, Bjorn Haave, Chelsea E. Cunningham, Yuliang Wu, Roland Arnold, Keith Bonham, Andrew Freywald, Jie Han, Franco J. Vizeacoumar

## Abstract

Can transcriptomic alterations drive the evolution of tumors? We rationalize that expressional changes found in all patients arise earlier in tumor development compared to alterations that occur only in limited subsets of patients. Our analyses of non-mutated genes from the non-amplified regions of the genome of 158 triple negative breast cancer (TNBC) cases identified 219 exclusively expression-altered (EEA) genes that may play important role in TNBC. Phylogenetic analyses of these genes predict a “punctuated burst” of multiple gene up-regulation events occurring at early stages of tumor development, followed by minimal subsequent changes later in tumor progression. Remarkably, this punctuated burst of expressional changes is instigated by hypoxia-related molecular events, predominantly in two groups of genes that control chromosomal instability (CIN) and remodel tumor microenvironment (TME). We conclude that alterations in the transcriptome are not stochastic and that early stage hypoxia induces CIN and TME remodeling to permit further tumor evolution.

## INTRODUCTION

The Darwinian model of clonal selection, where a subset of genetic lesions drives tumor evolution and progression in a step-wise manner ^1-4^, is widely accepted as the mode of evolution of malignant cells under therapy or basal conditions ^5-7^. However, recent findings in prostate, pancreatic or triple negative breast cancer (TNBC), challenge this paradigm and question if gradualism is indeed the single mode of evolution ^8-10^. It may be instead a punctuated burst of molecular alterations in the early stages of cancer, where changes in biological environment of growing tumors require massive adaptations in the molecular machinery of cancer cells ^11-13^. Usually, normal cells respond to stress by deploying repair or resistance tools to maintain their genetic integrity and assure survival ^14,15^. In contrast, cancer cells typically do not have intact repair tools, which lead to genetic instability. Chromosomal instability (CIN) is a form of genetic instability that causes changes in both the structure and number of chromosomes ^15-25^. For example, mutations in CIN genes like BRCA1/2 increase the number of deletions up to 50 bps, causing multiple defects within the genome ^26^. Progressive accumulation of CIN within a tumor allows development of cell populations with heterogeneous properties. Some of these cells will carry selective survival advantages and will be responsible for further tumor progression ^3^. Likewise, overexpression of APOBEC3, a member of the cytidine deaminase gene family, may generate frequent C>T base substitutions also leading to tumor heterogeneity and progression along the malignancy pathway ^27^. Understanding the sequence of molecular events essential for tumor evolution may not only benefit early detection of malignancies, but may also allow the development of more effective treatment and even prevention strategies. While the role of accumulating genetic mutations in tumor progression has been extensively discussed, it is still not clear how alterations in gene expression patterns contribute to tumor evolution.

Changes to gene expression can be brought about by number of factors, including epigenetic modifications, translation regulation, and differences in mRNA and protein stability_28_. For example, increased activities of growth factor, chemokine and cytokine receptors can set off specific signaling cascades and subsequent changes in gene expression, without any direct involvement of genetic mutations. However, what are the most significant changes that occur within the transcriptome of cancer cells and how they may contribute to tumor evolution is not clear. Here we use an aggressive malignancy, TNBC, as a model to explore the role of transcriptomic alterations during early stages that are caused not by genomic mutations, but exclusively by differential gene expression. We achieve this by focusing specifically on genes that are heavily up-regulated in the non-amplified regions of the genome. We focused specifically on up-regulated genes because direct inhibition of these molecules may provide viable cancer treatment/prevention options at early stages of tumor development. Remarkably, our analysis of RNA seq data in 158 TNBC cases revealed that there is indeed a punctuated burst of expressional changes in two major groups of genes controlled by hypoxia-related factors. These two groups included molecules that regulate CIN and remodel tumor microenvironment (TME). This not only reveals new potential targets for TNBC therapy, but also indicates a critical role for hypoxia in very early stages of tumor development.

## RESULTS

### A Multi-step process to identify differentially expressed genes in breast cancer

To identify genes with aberrant expression patterns, we initially curated all the genes that are differentially regulated. We used the breast cancer-specific data from The Cancer Genome Atlas (TCGA) that represents the largest collection of patient samples with information on the mutation status, copy number aberrations (CNA), as well as gene expression patterns at different stages of tumor development. Gene expression in breast tumor samples was compared to the expression of the matching genes in normal samples using fold change (FC) and false discovery rate (FDR) after Empirical Bayes moderated t-test with Benjamini-Hochberg correction. Genes were considered as up-regulated genes if FDR ≤ 0.01 and FC ≤ 2. Down-regulated genes were selected if FDR ≤ 0.01 and FC ≤ −2. Our initial analyses in overall breast cancer identified 586 genes that were up-regulated and 1446 genes that were down-regulated at multiple stages of cancer progression (Supplementary Fig. 1a). The overlap between all stages is presented in Supplementary Table 1a. We also ran a complementary analysis to identify differentially regulated genes in specifically in TNBC. We found 1127 genes to be up-regulated and 1752 genes down-regulated across multiple stages of TNBC (Fig. 1a). The overlap between all stages is shown in Supplementary Table 1b. The Gene Set Enrichment Analysis (GSEA) indicated that the up-regulated genes in TNBC are enriched for molecules involved in cell cycle regulation and chromatin organization (p<0.001) (Fig. 1b). Results of our GSEA analysis of genes differentially up-regulated in TNBC tumors correlated well with the previously reported, differentially regulated genes from an independent cohort (p<0.001) (Fig. 1b) ^29^, which provides an additional support for the relevance of our observations. While we found a higher abundance of down-regulated genes, compared to the up-regulated genes, no similar significant enrichment was observed within the pool of the down-regulated genes. Similar results were obtained for overall breast cancer (Supplementary Fig. 1b). Taken together, these observations indicate that the application of our approach to the analysis of TCGA data allows identifying subsets of genes differentially regulated in TNBC tumors.

**Figure 1:**
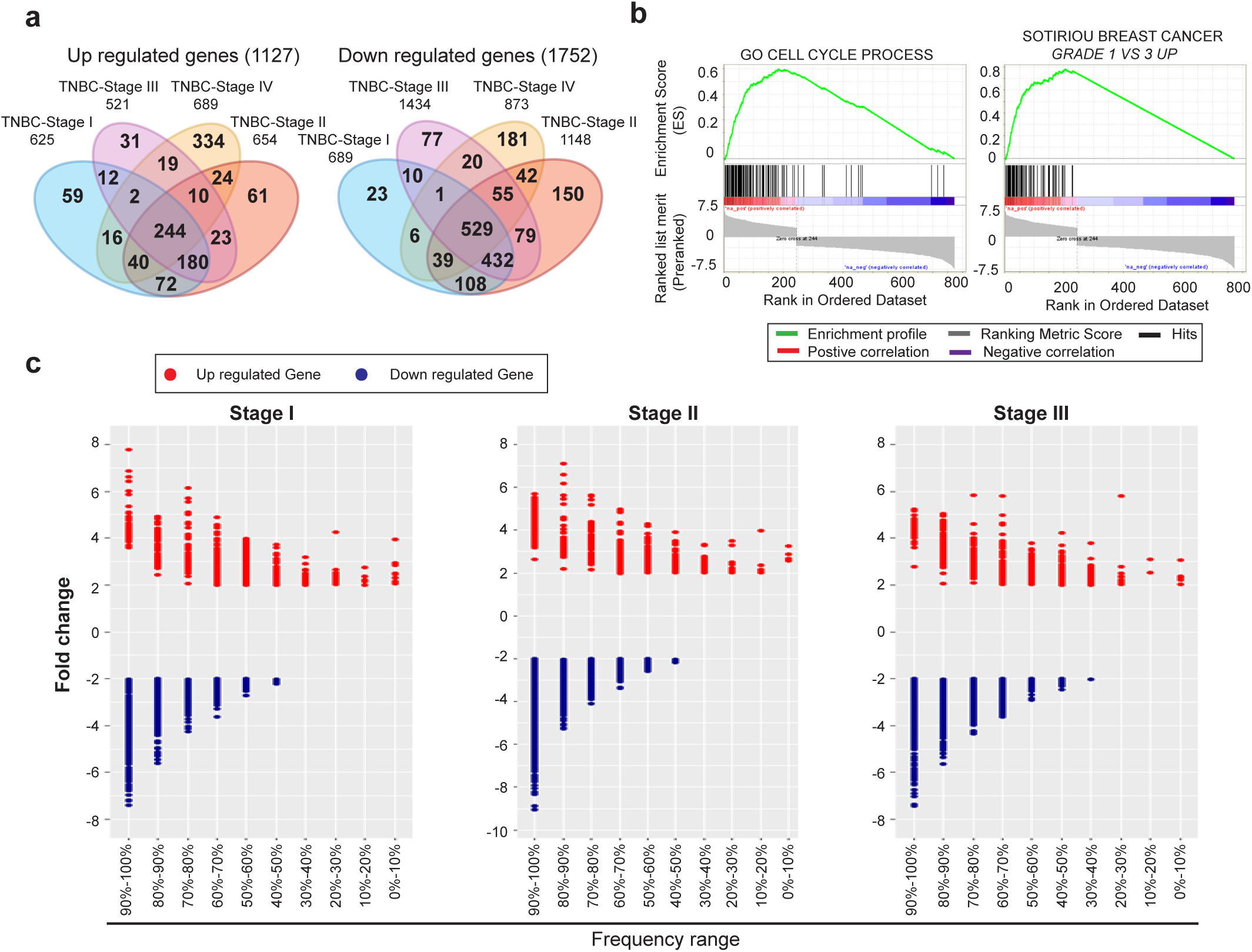
Identification of up-regulated genes in TNBC. **(a)** Venn diagram of differentially expressed genes in TNBC stage-specific tumors. The number of up and down-regulated genes at each stage of tumor and at the intersection between different stages have been represented. **(b)** Gene set enrichment analysis for up/down-regulated genes across all TNBC tumor stages. Gene Set Enrichment Analysis for 244 up-regulated genes (left) and 529 down-regulated genes (right) across four tumor stages along with previously identified, differentially up-regulated genes from Sotiriou et al ^29^**. (c)** Frequency distribution of differential expression in TNBC stage-specific tumors. Dot plot represents the fold change and the frequency range of TNBC stage-specific differentially expressed genes, where the red denotes up-regulated gene and the blue denotes down-regulated gene.

### Not all differentially expressed genes are equally deregulated across the population of breast cancer patients

While gene expression analysis to identify differentially regulated genes has been a common approach in cancer biology, we attempted to determine, how many of these genes are aberrantly expressed with high frequencies across the population of TNBC patients. We rationalized that common aberrations found in all patients should have arisen earlier in the development of the malignancy, compared to alterations that were found only in a subset of patients. Therefore, we have calculated a frequency of differential expression of each affected gene in TNBC tumors (Fig. 1c; Supplementary Fig. 1c). Throughout this analysis, we maintained a two-fold change in the expression level as a minimum requirement for a gene to be considered differentially regulated. The frequency of changes in each differentially expressed gene is calculated as a percentage of patients in whom the gene is up or down-regulated. We found 254 genes were up-regulated and 1197 genes were down-regulated in almost 70% of the TNBC patients (Supplementary Table 2a,b). Similar results for overall breast cancer are presented Supplementary Table 3a,b. Unfortunately, there were only two patient samples that were available in TNBC-stage IV in TCGA dataset, which was not sufficient to minimize random effects. Therefore, we computed our analyses using the larger number of samples involved in the first three stages of TNBC.

Changes in gene expression may not only arise from aberrant expression from an endogenous promoter, but also from accompanying chromosomal amplifications, deletions and other types of mutations. To account for this, we isolated the differentially regulated genes exclusively from the non-amplified/deleted regions of the genome. We identified 77 amplified chromosome regions from the TCGA dataset based on CNA, including several previously reported regions in 1q, 8q, 16p and 20q (Supplementary Table 4)^30^, as presented in the circos plot for TNBC (Fig. 2a) or overall breast cancer (Supplementary Fig. 2a). We further evaluated the concordance of amplification and gene expression by fold change with FDR and Pearson’s correlation. We considered genes likely to be driven by CNA if their Pearson’s correlation coefficient between expression and CNA was greater than 0.3, or they show significant differential CNA-associated expression change (Supplementary Fig. 2b,c; Supplementary Table 5a,b). Subsequently, we filtered out from our analysis 20 genes from TNBC patients that were in amplified regions or had strong correlations with chromosome amplification.

**Figure 2:**
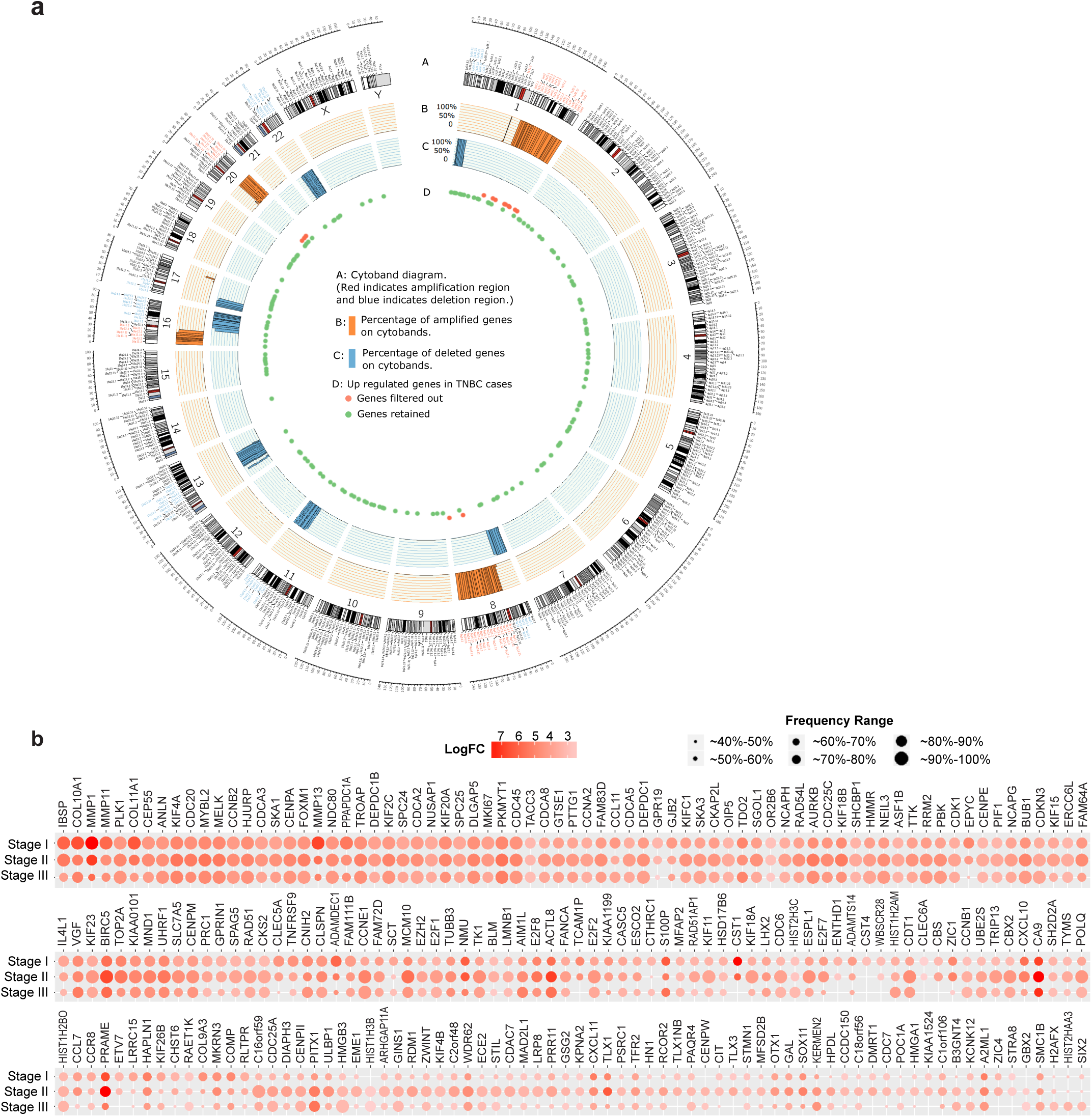
Elimination of amplified genes to identify 219 up-regulated events. **(a)** Amplified chromosome cytobands and up-regulated genes locus. Track A displays the cytoband diagram where the texts in red indicate identified amplified regions. Track B and C displays the frequency of genes showing amplification and deletion respectively in at least in 40% of patients in each cytoband. Genes in figure 1d were mapped to the Track D. **(b)** Fold change and frequency distribution for genes showing up-regulation in at least 70% of TNBC patients. Nodes in each column represent up-regulated genes with their sizes indicating the frequency of samples and their colors representing the fold change value in the specific tumor stage.

We also used somatic mutational analyses of 560 breast cancer whole genome sequencing database available at COSMIC to eliminate any gene that might be differentially expressed because of a mutation ^31^. By also excluding 13 genes whose loci information was ambiguous, we finally identified 219 exclusively expression-altered (EEA) genes that elevated their expression in TNBC (Fig. 2b) and therefore, may represent good therapeutic targets. Interestingly, we observed multiple distinct patterns of up-regulation with varying frequencies across different cancer stages (Fig. 2b). For example, some genes were constitutively up-regulated across all stages (PLK1, UBE2C or KIF4A). Similarly, certain genes were up-regulated mostly at later stages (CCNE1, HMGB3 or NUF2). In contrast to this category, some genes were up-regulated selectively at early stages but were gradually down-regulated through the later stages (MMP1, MMP11 or MMP13). Among the 219 up-regulation events, majority of changes occurred in chromosome 1 and 17 (Supplementary Fig. 2d). Surprisingly, although the expression of some initially up-regulated genes gradually decreased, we did not observe any instance where their expression returned back to normal levels (Supplementary Fig. 3). Importantly, while we find that not all up-regulated genes are overexpressed in all breast tumors across all cancer stages, our analysis has generated an explicit set of genes that are overexpressed in over 70% of patients at all stages of both all breast cancer and TNBC tumors (Supplementary Table 6a,b).

### Lineage analysis of up-regulation profiles reveals a punctuated pattern of evolution in early TNBC tumors

Previous studies have used somatic mutations and CNA to understand tumor evolution _8,10,11,13_. However, it is not clear, how alterations in gene expression may affect tumor progression. To further address this, we performed a clustering analysis on the expression profiles of the newly identified EEA genes to describe how they may influence TNBC progression. After identifying distinct tumor clusters, we constructed their distance tree to gain insights into their relationship, analogous to a phylogenetic analysis. We generated a binary matrix from profiles of the 219 EEA genes (0⁏= ⁏no up-regulation, 1⁏ = up-regulation) for each TNBC tumor. The pairwise Euclidian distances between samples were calculated and unsupervised hierarchical clustering with ward linkage^32^ was applied to identify homogenous groups, which allowed to cluster patients with similar up-regulation profiles. This approach (Fig. 3a) reveals that cluster C1 is diverged at the highest overhang with the highest dissimilarities from the remaining samples. In addition, several distinguishable branches C2, C3 and C4 are also clustered. The construction of the distance tree was based on the neighbor-joining algorithm^33^ to display the lineage between the four clusters. Assuming that the tumor is derived from a single or a group of homogenous normal cells and the complexity of up-regulation in a tumor increases with time, the history of its progression can be partially inferred by comparing homogeneous groups.

**Figure 3:**
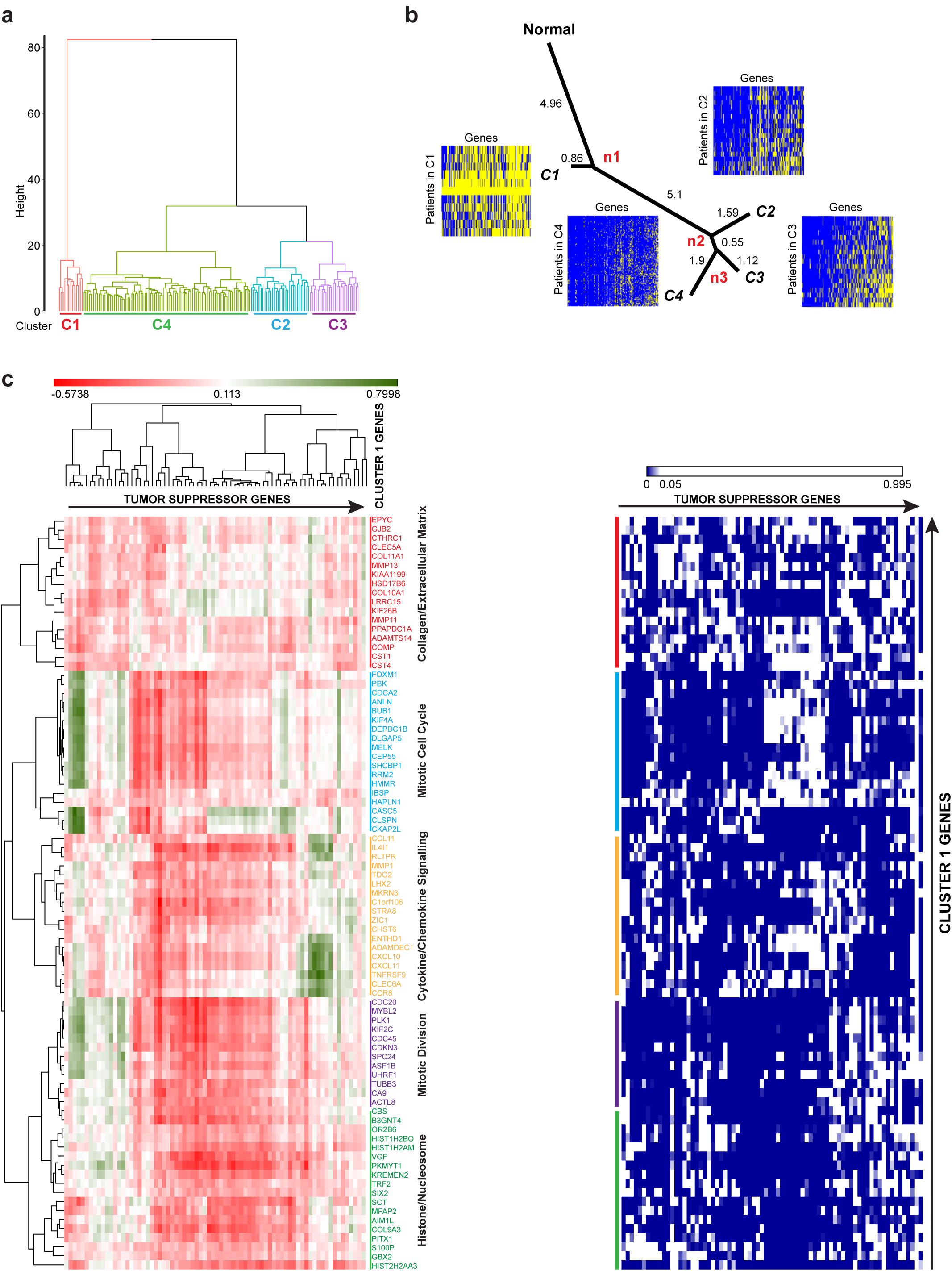
Identification of 83 up-regulation events that occur in early stages. **(a)** Hierarchical clustering of 219 EEA genes in TNBC patients. Different colors show TNBC patients clustered into four clusters represented as Red for Cluster 1 (C1), Purple for Cluster 2 (C2), Blue for Cluster 3 (C3) and Green for Cluster 4 (C4). **(b)** The progression of gene up-regulations in different TNBC clusters as shown by the phylogenetic tree. The figure shows the lineage of progression of gene up-regulations from the normal to distinct subpopulations. Heat maps with genes in columns and TNBC samples in rows display the up-regulation status (yellow: no up-regulation; blue: up-regulation) for different TNBC clusters. **(c)** Correlation clustergram of cluster 1 genes compared to known tumor suppressors. Red indicates negative correlation and green indicates positive correlation. The panel on the right represents, the significance of the correlation as a heat map. Blue indicates significance (<0.05) and white indicates lack of significance (>0.05).

The phylogenetic lineage showed that cluster C1 has the shortest distance from normal samples, which suggests that tumors in C1 may be recorded at the early tumor stage and the up-regulated genes in C1 may be of importance to tumor initiation and early progression (Fig. 3b). The lineage shows a large distance from normal cell to C1, indicating a large number of up-regulatory events are required for successful tumor progression through very early stages. C2, C3 and C4 clusters diverged for relatively small distances from the common ancestor n2, which suggests less dissimilarity from C2 to C3 and C4, indicating that minimal changes in gene expression were required at later stages. By measuring the Consine similarity between mean up-regulation profile and subset vector (See Methods for details), we identified the 83 EEA genes that may act as potential enabling factors within the early tumor evolution (Supplementary Table 7).

To confirm that the 83 up-regulation events are relevant to breast cancer progression, we next inquired if these changes in gene expression correlate with the loss of expression of known tumor suppressors. Vogelstein and colleagues identified ∼70 tumor suppressor genes that when inactivated by intragenic mutations can promote tumorigenesis^34^. We found a strong negative correlation in the expression of the 83 EEA genes and the 74 tumor suppressors (Fig. 3c). In summary, our analysis revealed that a large number of EEA events appear at the earliest stage of tumor development with fewer subsequent events at later stages, reflecting an emerging pressure from rapidly changing biological environment within early progressing tumors.

### TME remodeling and CIN cooperatively drive TNBC

Since our phylogenetic analysis indicated that the 83 up-regulated EEA genes are crucial early events in early tumorigenesis, we next explored the functionalities of these genes. Interestingly, we found a large subset of genes that are known to be involved in remodeling TME, including metalloproteinases (MMP1, MMP11, MMP13, ADAMDEC1, ADAMTS14), chemokine receptors and ligands (CXCL11, CXCL10, CCL11, CCR8), protease inhibitors (CST4, CST1), pH maintenance factors (CAIX), and different collagens (COL9A3, COL10A1). This emphasizes the critical role of extra-cellular matrix and TME remodeling at the early stage of tumor progression. Similarly, we also identified several of the EEA genes including, FOXM1, PLK1, BUB1, KIF2C, CDCA2, CDC20, CDKN3, KNL1 to name a few, that are known for their role in CIN and tumor development ^35-43^. This may reflect a selective pressure for additional genetic alterations in early tumors that would allow their further evolution. In addition, cluster 1 included genes like DEPDC1B and HMMR that have known roles in both TME remodeling as well as CIN associated functions ^44-48^. Overall, our identification of cluster 1 genes indicates that a punctuated burst of expressional changes occurs simultaneously in both CIN and TME remodeling genes very early in tumor development. Supplementary Table 8a lists literature evidence for the role of cluster 1 genes in TME and CIN.

If both CIN and TME remodeling ensue simultaneously, we should ask what possible factors could drive such punctuated burst. To address this, we used recently published causal analyses tools ^49^ available in the Ingenuity Pathway Analysis. In particular, we performed Upstream Regulator Analysis, and Causal Network Analysis to curate all interactions of cluster 1 genes (Fig. 4a,b). Interestingly, a large sub-set of direct up-stream interactions as well as causal interactions of both the CIN and TME genes (cluster 1), are hypoxia responsive genes ^50^ (Fig. 4a,b and Supplementary Table 8b,c). Invariably, almost 50% of the cluster 1 genes are also associated with poor prognosis (Fig.5a and Supplementary Fig. 4). This strongly suggests that very early in the course of tumor progression gradually increasing hypoxic conditions induce both CIN and TME remodeling to permit survival of cancer cells and their further evolution at later stages of malignancy.

**Figure 4:**
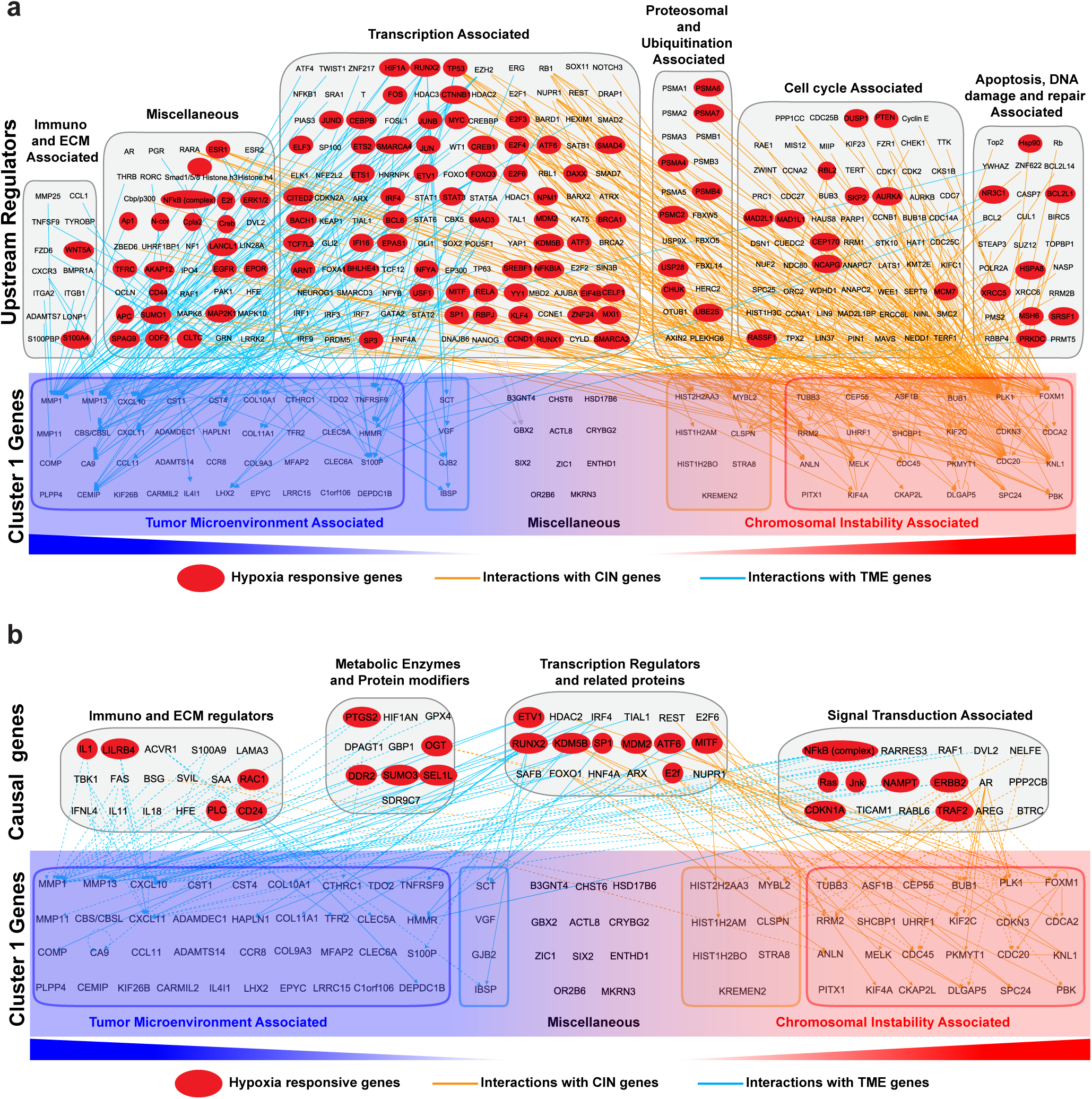
IPA analyses showing extensive interaction between hypoxia responsive genes with members of cluster 1 genes. **(a)** Upstream regulator analysis was performed with IPA for the cluster 1 genes and all the interactions retrieved are presented. Cluster 1 genes are classified into those that are associated with CIN or TME. The upstream genes that are hypoxia responsive, are highlighted in red. **(b)** Causal network analysis was performed with IPA for the cluster 1 genes and all the interactions retrieved are presented. Cluster 1 genes are classified into those that are associated with CIN or TME. The upstream genes that are hypoxia responsive, are highlighted in red.

Having identified a set of 83 EEA genes that act in early TNBC tumors, we sought to identify drugs that can benefit TNBC treatment at early stages and may potentially be also used for cancer prevention. To do this, we selected breast cancer cell lines that overexpress cluster 1 genes and analyzed their sensitivity to drugs using the cancerRXgene database (http://www.cancerrxgene.org). This database provides information on cell line drug sensitivity. The data for 265 drugs and multiple cell lines was examined to identify compounds that are more effective when used selectively with cell lines that highly express cluster 1 genes. We found four drugs, bleomycin, pevonedistat, ponatinib, and WIKI4, that showed a significant decrease in the IC50, for cell lines that highly expressed cluster 1 genes (Fig. 5b). Consistent with our identification of several cluster 1 genes being involved in CIN (Fig. 4a), our drug analyses indicate that cell lines with high expression of cluster 1 genes are more sensitive to a DNA damaging agent, bleomycin (Fig 5b).

**Figure 5:**
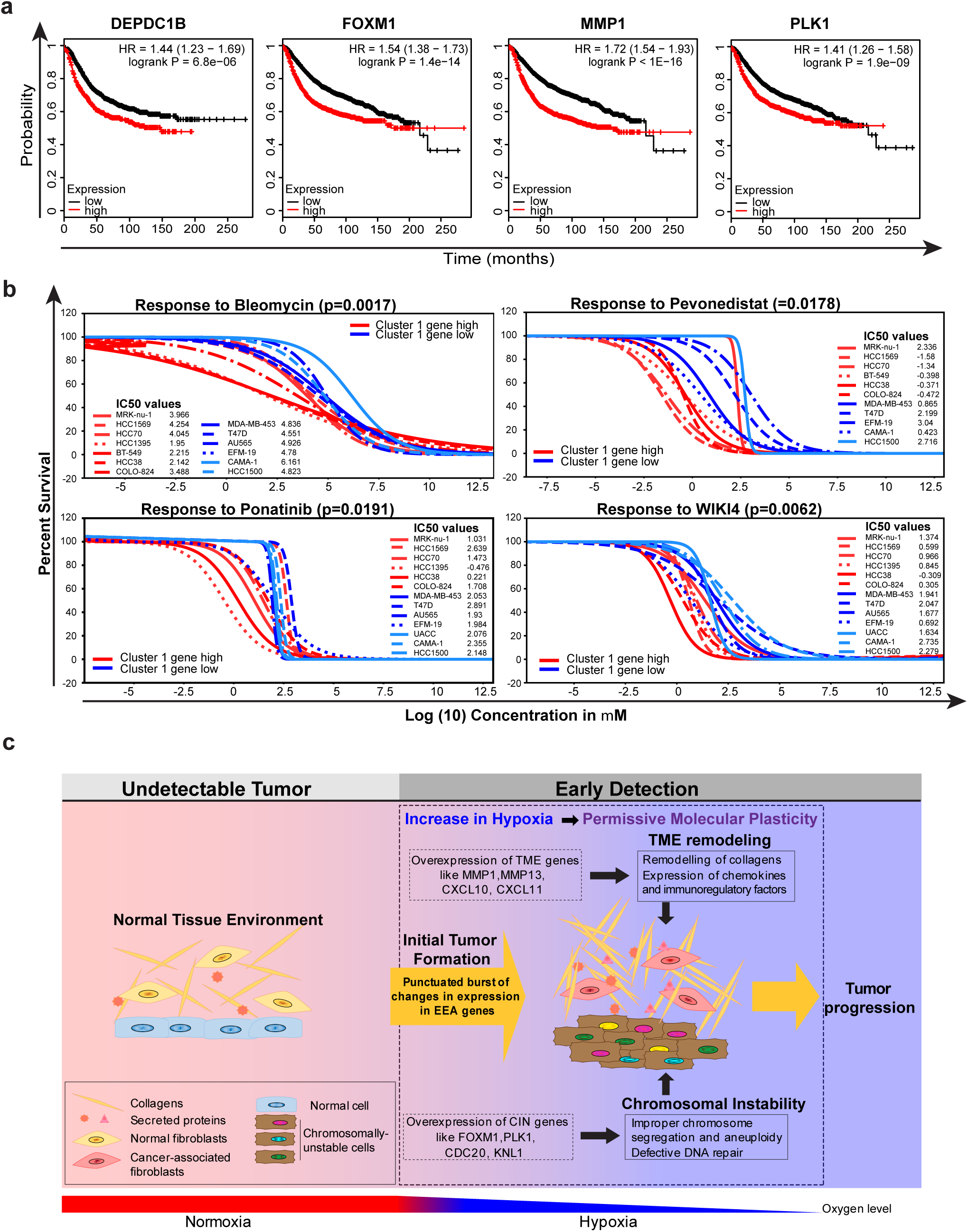
Survival plot, Drug response and a model describing the role of cluster 1 genes in tumor evolution. **(a)** Representative relapse free survival plots of breast cancer patients with low and high expression of cluster 1 genes **(b)** Dose response curves and IC50 values of drugs targeting cell lines with low and high expression of cluster 1 genes. **(c)** Schematic model showing the effect of simultaneous burst of CIN and TME-associated genes in response to hypoxia during early stages of cancer initiation.

## DISSCUSSION

Differential gene expression analyses have been traditionally used to examine fluctuations within the transcriptome in a given context for decades. This has been a powerful strategy to identify biomarkers and drug targets ^51-54^. However, tumor genome sequencing has provided new opportunities to re-examine these fluctuations in the context of tumor evolution. We rationalize that common aberrations detected across all patients arose earlier in the development of the malignancy compared to alterations that were found only in a subset of patients. Based on this, our strategy in this work is to explore the frequency of changes in the expression pattern of genes at different stages of TNBC progression. This is similar to previous studies that explored dynamic changes in mutations or CNA for a given patient at a given stage ^8-10,55,56^.

Changes in gene expression, unless constitutively observed, are often ignored as stochastic noise, specifically those that arise from variations in transcriptional regulation or biochemical modifications within cells. Our analyses deliver a number of important observations. First, compared to mutational changes, alterations within the transcriptome are more common and occur at high frequency. For example, the highly significant mutations in genes like PIK3CA or KRAS are observed in ∼30% of breast cancer patients. In contrast, overexpression of PLK1 or FOXM1 genes is observed in over 90% of patients (Supplementary Fig. 5). Second, more genes are down-regulated compared to up-regulated genes. Third, during tumor evolution, changes in expression pattern occur as punctuated bursts, where the initial singular burst results in simultaneous accumulation of overexpression of multiple EEA genes. Fourth, early changes in the expression of EEA molecules occur in genes that remodel TME and maintain chromosomal stability. This is most likely because survival within the progressively changing biological landscape during early stages requires cancer cells to both actively adjust to their microenvironment for their needs and to enhance CIN to facilitate their plasticity and adapt. Indeed, our unbiased genome-wide investigation reveals a strong functional connection between these two mechanisms and a crucial role of their coordinated effort in establishing early tumors. Interestingly, some TME genes, including MMP1, MMP11 and MMP13 proved to be up-regulated at early stages and gradually down-regulated through the later stages, although never achieving their normal levels. This suggests that their activities are essential at all stages of cancer progression, but their higher activity is required in early tumors, where the TME is not adjusted yet to the needs of malignant cells. Fifth, while we know that hypoxic TME can trigger tumor metastasis and invasion at later stages of cancer progression ^57,58^, our causal network analyses suggest that increasing hypoxia may be responsible for the cooperative induction of CIN and TME remodeling much earlier than previously appreciated (Fig. 5c). As hypoxic environment is also known to promote the propagation of tumor initiating cells (TICs) ^59,60^, we suspect that the expressional changes of EEA genes may facilitate this process. This is consistent with our finding that drugs like bleomycin and WIKI4 that efficiently eliminate TIC-enriched cell populations ^61,62^, cause selective lethality to cancer cell lines that overexpress cluster 1 genes (Fig. 5b).

Although, CIN is nearly ubiquitous in cancer cells, and is considered as an important factor in tumor development ^63^, our findings indicate that hypoxic TME of early tumor may function as a trigger of genetic instability. This model is consistent with previous observations, showing that repeated cycles of hypoxia, can down-regulate a number of DNA repair pathways in cancer cells, ultimately leading to genetic instability ^64,65^. In regards to this, the Glazer group has provided one of the first quantitative assessments of how genetic instability can be instigated by TME ^66^. Interestingly, several of the core EEA genes that maintain genome stability were experimentally shown to be involved in tumor development ^35-43^. Although some of these examples might be indicative of a direct role for CIN genes in tumorigenesis, in the context of our analyses, we suggest that overexpression of these genes may have enabled cancer cells to acquire properties that allowed them to survive at the early stage of cancer and thus, to develop detectable tumors (Fig.5c). In summary, our unbiased comprehensive analyses of the transcriptome directly link the early onset of hypoxia to the collective burst of CIN and TME remodeling factors in the initial stage of tumor progression, which highlights a therapeutic potential of targeting these molecules in TNBC tumors in their earliest detectable stage.

## METHODS

### Number of patient samples analyzed

We collected breast cancer samples from the Cancer Genome Atlas (TCGA) with information on the copy number aberration, gene expression as well as tumor information. According to the stage information, 1078 samples were classified into four tumor stages; from stage I to stage IV with tumors in stage V not being considered in this study. According to the IHC markers, 158 samples were classified as TNBC tumors in which the ER, PR and HER2 were all negative. With the tumor stage information, we classified TNBC tumors into TNBC-stage I, TNBC-stage II, TNBC-stage III and TNBC-stage IV. Similarly, TNBC tumors in stage IV were excluded. Moreover, 114 normal samples were collected from TCGA for comparison with tumor sample data. The numbers of samples in each stage and TNBC-stage specific samples are displayed in the Supplemental Table 9.

### Fold-change and FDR calculation

We applied two criteria, fold change and FDR calculation on the selection of differentially expressed genes. Fold-change is a biological assessment of changes in gene expression that is estimated by log2 (ratio), as represented in Equation 1, where the ratio of average expression of gene *i* in patients to the average expression in normal samples is calculated.

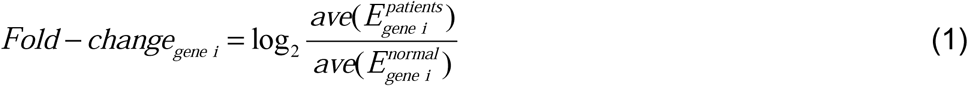

Empirical Bayes moderated t-test was applied to assess the statistical significance of differential expression. False discover rate (FDR) was obtained after Benjamini and Hochberg correction. We employed the Limma package ^67^ to derive the two assessments of differentially expressed genes.

### Computing frequency of differential expression in stage-specific patients

After the identification of up/down-regulated genes in each stage and TNBC-stage patients, next we aimed to evaluate the frequency of identified differential expression in stage-specific patients. For each tumor stage, we calculated the fold change of identified up or down-regulated gene *I* by comparing the expression in patient *j* to the average expression in normal samples (Equation 2). Following this, the frequency of patients in which the fold-change of gene *i* is greater than 2 or less than −2 was calculated.

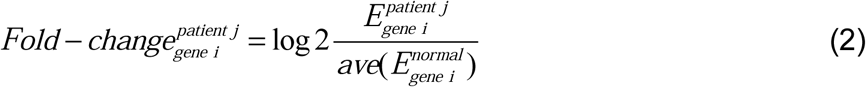

### Evaluating the concordance between copy number amplification and up-regulated gene expression

As changes in gene expression may arise from the chromosomal amplification, here we aimed to evaluate the associations between copy number amplification (CNA) and up-regulations in gene expression and identify the up-regulations that are driven by CNA. We generated the CNA profile with patients (in rows) and genes (in columns) from the data obtained from TCGA. Only the data of patients whose CNA profile and up-regulation status are available were considered for this study. Using a scoring system, genes getting amplified in a patient were represented as 1 or otherwise 0 and the amplified genes were grouped based on the profile scoring. To avoid patient heterogeneities, only genes showing amplification over 40% of patients were considered as cancer relevant amplified genes. CNA regions were identified by calculating the percentage of amplification of genes on each chromosome region, and regions with at least 40% of amplification genes were identified as CNA regions. Then we analyzed the concordance between CNA and up-regulations in gene expression by two evaluation ways. Primarily, cfor each up-regulated gene *i* at CNA regions, according to the gene *i*’s amplification status, the patient set *S* were grouped into two sets 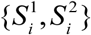, where 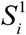 and 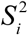 denotes patients with and without gene *i* getting amplified respectively. The fold CNA-associated change was calculated by comparing the difference between the log2 of mean expression of gene *i* in 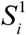 and 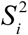. Meanwhile empirical bayes moderated t-test is applied on the two groups. To correct for multiple hypothesis testing, the p value was converted to FDR by Benjamini-Hochberg correction. If gene *i* showed at least 1 positive fold CNA-associated change and FDR smaller than 0.01, it is considered to be associated strongly with CNA. Secondly, Pearson’s correlation coefficient was calculated to quantify the correlation between CNA and gene expression. If a gene showed the value of Pearson’s correlation coefficient larger than 0.3, it is considered as CNA-driven genes as well.

### TNBC subpopulations and neighboring-joining algorithm analyses

To identify the distinguishable TNBC subpopulations, which reveal similar up-regulation profiles, we performed clustering analysis within TNBC patients. The binary matrix was generated with up-regulation status in rows and patient in columns. If a gene is up-regulated in a patient, it was indicated by 1, otherwise 0 using the scoring system described in the previous section. Then we calculated pairwise Euclidian distance between patients and performed hierarchical clustering that clustered TNBC patients into distinguishable clusters. The mean up-regulation profile for each cluster was generated to represent the up-regulation status. Assuming that no gene up-regulations appeared before the initiation of cancer, the profile with all zeros was generated to represent normal samples. The distance tree was constructed based on profiles in normal and mean profiles by Euclidian distance and neighbor-joining algorithm.

### Assigning genes into different subpopulations

To determine the appearance of up-regulations in various subsets of the four subpopulations, we generated the four-dimensional binary vectors to represent each of the fifteen possible subsets of the four subpopulations, from (0, 0, 0, 1) to (1, 1, 1, 1). Four each gene*i*, the consine similarities between mean profile Equation.3. *p_i_* and each subset vector *v_j_* is calculated by the

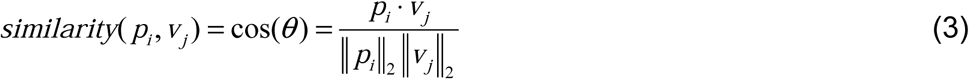

The gene was assigned to the subset vector with the maximum similarity to its mean profile^68^. For example, if a gene has the maximum similarity with the subset vector (0, 1, 1, 1), it means it getting up-regulated in subpopulations 2, 3 and 4.

### IPA and hypoxia analysis

IPA analysis was performed on the genes from cluster 1 as described in Kramer et al.,^49^. The gene list was first annotated and the data set underwent various analyses including for core expression to study the interactions. The gene interactions were explored, built and different overlays including pathways, disease and function and molecule activity prediction were applied to obtain the required outputs. Comparison analysis was also performed among the different subpopulations (referred as clusters). Hypoxia analyses were performed using the hypoxia database (http://www.hypoxiadb.com). This database includes 72,000 manually curated entries taken on 3500 proteins extracted from 73 peer-reviewed publications selected from PubMed. As described in Khurana et al., it provides manually curated literature references to support the inclusion of the protein in the database and establish its association with hypoxia ^50^.

### Drug Data analyses

The cell lines from the cancerRXgene database were divided into high cluster 1 expression and low cluster 1 expression. This was done by creating a table where the rows were cell lines, the columns were cluster 1 genes and the intersection at each row and column was the expression value of that gene in that cell line. The expression values across all cell lines for each gene were then added together and the mean and standard deviation were calculated. Then each cell line was given a Z score for that gene 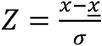 where x is the value, <u>x</u> is the mean and *σ* is the standard deviation. Each cell line then had all of its Z scores summed together to give the total score of c1 gene expression. The 85th percentile and the 15th percentile were then taken to be the cluster 1 high expression and cluster 1 low expression groups respectively. After grouping the cell lines into high and low expression of cluster 1, analysis was ran on the sensitivity of these cell lines to drugs. The drug data was obtained from the cancerrxgene database. For each drug, the IC50 values from the database were taken for each cell line of the high expression cell lines, and each of the low expression cell lines. The IC50 values for each group were then compared using a Mann-Whitney-U test. Using the percent survival, we generated the graphs and dose data from the cancerrxgene database and fitting a sigmoidal curve to the resulting plot, using the literature IC50 value as an estimator. The sigmoid curve used was of the form 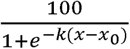

## Acknowledgements

We thank members of the Vizeacoumar, Han and Freywald laboratories for their feedback and comments. C.E.C. is supported by Lisa Rendall Breast Cancer fellowship. S.P. is supported by SCA post-doctoral fellowship.

## List of Abbreviations

CIN: chromosomal instability
TNBC: triple negative breast cancer
TCGA: The Cancer Genome Atlas
CNV: copy number variation

## Supplementary Figures

**Supplementary Figure 1: Identification of up-regulated genes in overall breast cancer.**

**(a)** Venn diagram of differentially expressed genes in overall (top) and ER+ patients (bottom) respectively. **(b)** Gene set enrichment analysis for up/down-regulated genes across all tumor stages. It shows the Gene Set Enrichment Analysis for 307 up-regulated genes (left) and 922 down-regulated genes (right) across four tumor stages along with previously identified, differentially up-regulated genes from Sotiriou et al ^29^**. (c)** Frequency distribution of differential expression in overall patients. This plot represents the fold change and the frequency range of stage-specific differentially expressed genes, where the red denotes up-regulated gene and the blue denotes down-regulated gene.

**Supplementary Figure 2: Expression pattern of up-regulated genes. (a)** Amplified chromosome cytobands and up-regulated genes locus. Track A displays the cytoband diagram where the texts in red indicate identified amplified regions. Track B and C displays the frequency of genes showing amplification and deletion respectively at least in 40% of patients in each cytoband. Genes in supplementary figure 2a were mapped to the Track D. **(b and c)** The evaluation on the concordance between gene expression and amplification. Nodes represent up-regulated genes in overall breast cancer cases in amplified regions, showing Pearson’s correlation coefficient and fold CNA-associated change. Genes in red were considered driven by copy number amplification either in overall breast cancer (B) or in TNBC (C). **(d)** Distribution of the 219 up-regulated events according to their chromosomal location.

**Supplementary Figure 3:** Box plots of Cluster 1 gene expression at various stages of TNBC tumor. The y-axis represents log2-transformed gene expression and x-axis denotes TNBC stages.

**Supplementary Figure 4:** Relapse free survival plot in breast cancer patients having low and high expression of cluster 1 genes.

**Supplementary Figure 5:** Percentage of patients that either overexpress or carry mutations in some of the key cancer genes.

## Supplementary Tables

**Supplementary Table 1:** Up/down-regulated genes across all stages in overall patients, TNBC patients and ER+ patients, respectively. Table 1a, and 1b show up/down-regulated genes in different tumor stages for overall and TNBC cases, respectively.

**Supplementary Table 2:** Stage-specific up/down regulated genes in at least 70% of TNBC patients. Table 2a and 2b show up and down regulated genes in at least 70% of TNBC patients in each stage, respectively, as well as the fold change, frequency and FDR information.

**Supplementary Table 3:** TNBC stage-specific differentially expressed genes in at least 70% of overall patients. Table 3a and 3b show up and down regulated genes in at least 70% of overall patients in each stage, respectively, as well as the fold change, frequency and FDR information.

**Supplementary Table 4:** Identified amplified regions. Table 4 shows the identified chromosomal amplification regions.

**Supplementary Table 5:** Evaluations on the concordances between gene expression and copy number amplification for up-regulated genes on amplified regions. Table 5a and 5b show Pearson’s correlation coefficient between gene expression and CNA, and fold CNA-associated change with FDR, and cytoband for up-regulated genes in overall and TNBC patients, respectively.

**Supplementary Table 6:** EEA genes in overall and TNBC patients. Table 6a and 6b show EEA genes in overall and TNBC patients, respectively.

**Supplementary Table 7:** Gene lists in four clusters. Table 7a, 7b, 7c and 7d show genes in cluster 1, 2, 3 and 4, respectively.

**Supplementary Table 8:** Analyses of cluster 1 genes and its interactions. Table 8a lists the cluster 1 genes and their association with CIN or TME. Table 8b and 8c enlist the upstream regulator and causal network genes respectively, along with their association with CIN or TME and hypoxia responsiveness.

**Supplementary Table 9:** The number of genes in each tumor stage and subtype. Table 8 shows the number of ER+, HER2+, TNBC, and overall patients in different stages.

## REFERENCES

1. Burrell, R. A., McGranahan, N., Bartek, J. & Swanton, C. The causes and consequences of genetic heterogeneity in cancer evolution. Nature 501, 338–345, doi:10.1038/nature12625 (2013).

2. Burrell, R. A. & Swanton, C. The evolution of the unstable cancer genome. Current opinion in genetics & development 24, 61–67, doi:10.1016/j.gde.2013.11.011 (2014).

3. McGranahan, N. & Swanton, C. Biological and Therapeutic Impact of Intratumor Heterogeneity in Cancer Evolution. Cancer Cell 27, 15–26, doi:10.1016/j.ccell.2014.12.001 (2015).

4. Swanton, C. Intratumor heterogeneity: evolution through space and time. Cancer Res 72, 4875–4882, doi:10.1158/0008-5472.CAN-12-2217 (2012).

5. Armitage, P. & Doll, R. The age distribution of cancer and a multi-stage theory of carcinogenesis. British journal of cancer 8, 1–12 (1954).

6. Nordling, C. O. A new theory on cancer-inducing mechanism. British journal of cancer 7, 68–72 (1953).

7. Nowell, P. C. The clonal evolution of tumor cell populations. science 194, 23–28 (1976).

8. Gao, R. et al. Punctuated copy number evolution and clonal stasis in triple-negative breast cancer. Nat Genet 48, 1119–1130, doi:10.1038/ng.3641 (2016).

9. Notta, F. et al. A renewed model of pancreatic cancer evolution based on genomic rearrangement patterns. Nature 538, 378–382, doi:10.1038/nature19823 (2016).

10. Baca, S. C. et al. Punctuated evolution of prostate cancer genomes. Cell 153, 666–677, doi:10.1016/j.cell.2013.03.021 (2013).

11. Genomic Analysis of Pancreatic Cancers Reveals Punctuated Evolution. Cancer Discov 6, OF17, doi:10.1158/2159-8290.CD-RW2016-198 (2016).

12. Cross, W., Graham, T. A. & Wright, N. A. New paradigms in clonal evolution: punctuated equilibrium in cancer. The Journal of pathology 240, 126–136, doi:10.1002/path.4757 (2016).

13. Davis, A., Gao, R. & Navin, N. Tumor evolution: Linear, branching, neutral or punctuated? Biochim Biophys Acta 1867, 151–161, doi:10.1016/j.bbcan.2017.01.003 (2017).

14. Jallepalli, P. V. & Lengauer, C. Chromosome segregation and cancer: cutting through the mystery. Nat Rev Cancer 1, 109–117, doi:10.1038/35101065 (2001).

15. Nowak, M. A. et al. The role of chromosomal instability in tumor initiation. Proc Natl Acad Sci U S A 99, 16226–16231, doi:10.1073/pnas.202617399 (2002).

16. Roschke, A. V. & Rozenblum, E. Multi-layered cancer chromosomal instability phenotype. Frontiers in oncology 3, 302, doi:10.3389/fonc.2013.00302 (2013).

17. Schvartzman, J. M., Sotillo, R. & Benezra, R. Mitotic chromosomal instability and cancer: mouse modelling of the human disease. Nat Rev Cancer 10, 102–115, doi:10.1038/nrc2781 (2010).

18. Holland, A. J. & Cleveland, D. W. Boveri revisited: chromosomal instability, aneuploidy and tumorigenesis. Nat Rev Mol Cell Biol 10, 478–487, doi:10.1038/nrm2718 (2009).

19. Thompson, S. L. & Compton, D. A. Examining the link between chromosomal instability and aneuploidy in human cells. J Cell Biol 180, 665–672, doi:10.1083/jcb.200712029 (2008).

20. Geigl, J. B., Obenauf, A. C., Schwarzbraun, T. & Speicher, M. R. Defining ‘chromosomal instability’. Trends Genet 24, 64–69, doi:10.1016/j.tig.2007.11.006 (2008).

21. Barber, T. D. et al. Chromatid cohesion defects may underlie chromosome instability in human colorectal cancers. Proc Natl Acad Sci U S A 105, 3443–3448, doi:10.1073/pnas.0712384105 (2008).

22. Weaver, B. A. & Cleveland, D. W. Does aneuploidy cause cancer? Current opinion in cell biology 18, 658–667, doi:10.1016/j.ceb.2006.10.002 (2006).

23. Gollin, S. M. Mechanisms leading to chromosomal instability. Semin Cancer Biol 15, 33–42, doi:10.1016/j.semcancer.2004.09.004 (2005).

24. Grigorova, M., Staines, J. M., Ozdag, H., Caldas, C. & Edwards, P. A. Possible causes of chromosome instability: comparison of chromosomal abnormalities in cancer cell lines with mutations in BRCA1, BRCA2, CHK2 and BUB1. Cytogenetic and genome research 104, 333–340, doi:10.1159/000077512 (2004).

25. Sieber, O. M., Heinimann, K. & Tomlinson, I. P. Genomic instability-the engine of tumorigenesis? Nat Rev Cancer 3, 701–708, doi:10.1038/nrc1170 (2003).

26. Alexandrov, L. B. et al. Signatures of mutational processes in human cancer. Nature 500, 415–421, doi:10.1038/nature12477 (2013).

27. Harris, R. S. Molecular mechanism and clinical impact of APOBEC3B-catalyzed mutagenesis in breast cancer. Breast cancer research : BCR 17, 8, doi:10.1186/s13058-014-0498-3 (2015).

28. Arcondeguy, T., Lacazette, E., Millevoi, S., Prats, H. & Touriol, C. VEGF-A mRNA processing, stability and translation: a paradigm for intricate regulation of gene expression at the post-transcriptional level. Nucleic Acids Res 41, 7997–8010, doi:10.1093/nar/gkt539 (2013).

29. Sotiriou, C. et al. Gene expression profiling in breast cancer: understanding the molecular basis of histologic grade to improve prognosis. Journal of the National Cancer Institute 98, 262–272, doi:10.1093/jnci/djj052 (2006).

30. Guenthoer, J. et al. Assessment of palindromes as platforms for DNA amplification in breast cancer. Genome Res 22, 232–245, doi:10.1101/gr.117226.110 (2012).

31. Nik-Zainal, S. et al. Landscape of somatic mutations in 560 breast cancer whole-genome sequences. Nature 534, 47–54, doi:10.1038/nature17676 (2016).

32. Johnson, S. C. Hierarchical clustering schemes. Psychometrika 32, 241–254 (1967).

33. Saitou, N. & Nei, M. The neighbor-joining method: a new method for reconstructing phylogenetic trees. Molecular biology and evolution 4, 406–425 (1987).

34. Wood, L. D. et al. The genomic landscapes of human breast and colorectal cancers. science 318, 1108–1113, doi:10.1126/science.1145720 (2007).

35. Wang, I. C. et al. Forkhead box M1 regulates the transcriptional network of genes essential for mitotic progression and genes encoding the SCF (Skp2-Cks1) ubiquitin ligase. Molecular and cellular biology 25, 10875–10894, doi:10.1128/MCB.25.24.10875-10894.2005 (2005).

36. Yu, G. et al. FoxM1 promotes breast tumorigenesis by activating PDGF-A and forming a positive feedback loop with the PDGF/AKT signaling pathway. Oncotarget 6, 11281–11294, doi:10.18632/oncotarget.3596 (2015).

37. Smith, M. R. et al. Malignant transformation of mammalian cells initiated by constitutive expression of the polo-like kinase. Biochem Biophys Res Commun 234, 397–405 (1997).

38. Li, Z. et al. Polo-like kinase 1 (Plk1) overexpression enhances ionizing radiation-induced cancer formation in mice. The Journal of biological chemistry, doi:10.1074/jbc.M117.810960 (2017).

39. Ricke, R. M., Jeganathan, K. B. & van Deursen, J. M. Bub1 overexpression induces aneuploidy and tumor formation through Aurora B kinase hyperactivation. J Cell Biol 193, 1049–1064, doi:10.1083/jcb.201012035 (2011).

40. de Carcer, G. & Malumbres, M. A centrosomal route for cancer genome instability. Nat Cell Biol 16, 504–506, doi:10.1038/ncb2978 (2014).

41. Nam, H. J. & van Deursen, J. M. Cyclin B2 and p53 control proper timing of centrosome separation. Nat Cell Biol 16, 538–549, doi:10.1038/ncb2952 (2014).

42. Li, M., Fang, X., Wei, Z., York, J. P. & Zhang, P. Loss of spindle assembly checkpoint-mediated inhibition of Cdc20 promotes tumorigenesis in mice. J Cell Biol 185, 983–994, doi:10.1083/jcb.200904020 (2009).

43. Urata, Y. N., Takeshita, F., Tanaka, H., Ochiya, T. & Takimoto, M. Targeted Knockdown of the Kinetochore Protein D40/Knl-1 Inhibits Human Cancer in a p53 Status-Independent Manner. Scientific reports 5, 13676, doi:10.1038/srep13676 (2015).

44. Marchesi, S. et al. DEPDC1B coordinates de-adhesion events and cell-cycle progression at mitosis. Dev Cell 31, 420–433, doi:10.1016/j.devcel.2014.09.009 (2014).

45. Yang, Y. et al. DEPDC1B enhances migration and invasion of non-small cell lung cancer cells via activating Wnt/beta-catenin signaling. Biochem Biophys Res Commun 450, 899–905, doi:10.1016/j.bbrc.2014.06.076 (2014).

46. Chen, D. et al. Phosphorylation of DEPDC1 at Ser110 is required to maintain centrosome organization during mitosis. Experimental cell research 358, 101–110, doi:10.1016/j.yexcr.2017.06.005 (2017).

47. Connell, M. et al. HMMR acts in the PLK1-dependent spindle positioning pathway and supports neural development. eLife 6, doi:10.7554/eLife.28672 (2017).

48. Veiseh, M. & Turley, E. A. Hyaluronan metabolism in remodeling extracellular matrix: probes for imaging and therapy of breast cancer. Integr Biol (Camb) 3, 304–315, doi:10.1039/c0ib00096e (2011).

49. Kramer, A., Green, J., Pollard, J., Jr. & Tugendreich, S. Causal analysis approaches in Ingenuity Pathway Analysis. Bioinformatics 30, 523–530, doi:10.1093/bioinformatics/btt703 (2014).

50. Khurana, P., Sugadev, R., Jain, J. & Singh, S. B. HypoxiaDB: a database of hypoxia-regulated proteins. Database (Oxford) 2013, bat074, doi:10.1093/database/bat074 (2013).

51. Glas, A. M. et al. Converting a breast cancer microarray signature into a high-throughput diagnostic test. BMC Genomics 7, 278, doi:10.1186/1471-2164-7-278 (2006).

52. Ludwig, J. A. & Weinstein, J. N. Biomarkers in cancer staging, prognosis and treatment selection. Nat Rev Cancer 5, 845–856, doi:10.1038/nrc1739 (2005).

53. Scherf, U. et al. A gene expression database for the molecular pharmacology of cancer. Nat Genet 24, 236–244, doi:10.1038/73439 (2000).

54. Frank, R. & Hargreaves, R. Clinical biomarkers in drug discovery and development. Nat Rev Drug Discov 2, 566–580, doi:10.1038/nrd1130 (2003).

55. Sun, Y., Yao, J., Nowak, N. J. & Goodison, S. Cancer progression modeling using static sample data. Genome biology 15, 440, doi:10.1186/s13059-014-0440-0 (2014).

56. Navin, N. et al. Inferring tumor progression from genomic heterogeneity. Genome Res 20, 68–80, doi:10.1101/gr.099622.109 (2010).

57. Rankin, E. B. & Giaccia, A. J. Hypoxic control of metastasis. science 352, 175–180, doi:10.1126/science.aaf4405 (2016).

58. Schito, L. & Rey, S. Hypoxic pathobiology of breast cancer metastasis. Biochim Biophys Acta 1868, 239–245, doi:10.1016/j.bbcan.2017.05.004 (2017).

59. Lin, Q. & Yun, Z. Impact of the hypoxic tumor microenvironment on the regulation of cancer stem cell characteristics. Cancer biology & therapy 9, 949–956 (2010).

60. Carnero, A. & Lleonart, M. The hypoxic microenvironment: A determinant of cancer stem cell evolution. Bioessays 38 Suppl 1, S65–74, doi:10.1002/bies.201670911 (2016).

61. Nicolay, N. H. et al. Mesenchymal stem cells are sensitive to bleomycin treatment. Scientific reports 6, 26645, doi:10.1038/srep26645 (2016).

62. James, R. G. et al. WIKI4, a novel inhibitor of tankyrase and Wnt/ss-catenin signaling. PLoS One 7, e50457, doi:10.1371/journal.pone.0050457 (2012).

63. Hanahan, D. & Weinberg, R. A. Hallmarks of cancer: the next generation. Cell 144, 646–674, doi:10.1016/j.cell.2011.02.013 (2011).

64. Klein, T. J. & Glazer, P. M. The tumor microenvironment and DNA repair. Semin Radiat Oncol 20, 282–287, doi:10.1016/j.semradonc.2010.05.006 (2010).

65. Yuan, J., Narayanan, L., Rockwell, S. & Glazer, P. M. Diminished DNA repair and elevated mutagenesis in mammalian cells exposed to hypoxia and low pH. Cancer Res 60, 4372– 4376 (2000).

66. Reynolds, T. Y., Rockwell, S. & Glazer, P. M. Genetic instability induced by the tumor microenvironment. Cancer Res 56, 5754–5757 (1996).

67. Ritchie, M. E. et al. limma powers differential expression analyses for RNA-sequencing and microarray studies. Nucleic Acids Res 43, e47, doi:10.1093/nar/gkv007 (2015).

68. Navin, N. et al. Tumour evolution inferred by single-cell sequencing. Nature 472, 90–94, doi:10.1038/nature09807 (2011).

